# Uncovering Differences in Hydration Free Energies and Structures for Model Compound Mimics of Charged Sidechains of Amino Acids

**DOI:** 10.1101/2021.02.04.429838

**Authors:** Martin J. Fossat, Xiangze Zeng, Rohit V. Pappu

**Affiliations:** Department of Biomedical Engineering and Center for Science & Engineering of Living Systems (CSELS) Washington University in St. Louis, St. Louis, MO 63130, USA

## Abstract

Free energies of hydration are of fundamental interest for modeling and understanding conformational and phase equilibria of macromolecular solutes in aqueous phases. Of particular relevance to systems such as intrinsically disordered proteins are the free energies of hydration and hydration structures of model compounds that mimic charged sidechains of Arg, Lys, Asp, and Glu. Here, we deploy a Thermodynamic Cycle based Proton Dissociation (TCPD) approach in conjunction with data from direct measurements to obtain estimates for the free energies of hydration for model compounds that mimic the sidechains of Arg^+^, Lys^+^, Asp^-^, and Glu^-^. Irrespective of the choice made for the hydration free energy of the proton, the TCPD approach reveals clear trends regarding the free energies of hydration for Arg^+^, Lys^+^, Asp^-^, and Glu^-^. These trends include asymmetries between the hydration free energies of acidic (Asp^-^ and Glu^-^) and basic (Arg^+^ and Lys^+^) residues. Further, the TCPD analysis, which relies on a combination of experimental data, shows that the free energy of hydration of Arg^+^ is less favorable than that of Lys^+^. We sought a physical explanation for the TCPD derived trends free energies of hydration. To this end, we performed temperature dependent calculations of free energies of hydration and analyzed hydration structures from simulations that use the polarizable AMOEBA (Atomic Multipole Optimized Energetics for Biomolecular Applications) forcefield and water model. At 298 K, the AMOEBA model generates estimates of free energies of hydration that are consistent with TCPD values with a free energy of hydration for the proton of ≈ -259 kcal / mol. Analysis of temperature dependent simulations leads to a structural explanation for the observed differences in free energies of hydration of ionizable residues and reveals that the heat capacity of hydration is positive for Arg^+^ and Lys^+^ and negative for Asp^-^ and Glu^-^.

## 1. INTRODUCTION

There is growing interest in uncovering the sequence-specific conformational preferences of intrinsically disordered proteins (IDPs)^1^, and in using these insights to quantify sequence-specific contributions to the driving forces for phase separation ^2^. In a purely additive model ^3^, sequence-ensemble relationships of IDPs can be rationalized using the free energies of hydration of model compound mimics of sidechain and backbone moieties ^4^. Indeed, the molecular transfer model of Thirumalai coworkers^5^ is a direct illustration of how free energies of solvation can be used to obtain predictive, coarse-grained descriptions of conformational transitions of proteins as a function of changes to solution conditions ^6^.

At a specific temperature and pressure, the free energy of hydration (Δµ_h_) is defined as the change in free energy associated with transferring the solute of interest from a dilute vapor phase into water ^7^. Vapor pressure osmometry was an early method adopted by Wolfenden^8^ to measure free energies of hydration. While this works for polar solutes, including neutral forms of ionizable species, it cannot be used to measure free energies of hydration of ionizable residues because of the ultra-low vapor pressures and the confounding effects of ion-pairing in the gas phase. Calorimetry ^9^ is also problematic because of the large magnitudes of free energies of hydration for ionizable residues ^10^. And because stable solutions are electroneutral ^10b^, estimates of free energies of hydration of ionic species have to rely on parsing numbers derived from measurements of whole salts against those of a suitable reference system ^11^. Parsing measurements for whole salts also rests on the assumption of minimal ion-pairing or clustering, which need not be true in general, especially for organic ions ^12^.

Here, we incorporate updated estimates for a series of experimentally measured quantities and combine these with a **T**hermodynamic **C**ycle based on **P**roton **D**issociation (TCPD) – see **Figure 1** – to obtain a distribution of experimentally derived estimates for free energies of hydration of charged amino acids at 298 K ^13^. This approach uses inputs from: (i) direct measurements of proton dissociation energies in the gas phase; (ii) measured pK_a_ values in the aqueous phase – including recent updates based on revisited measurements for the pK_a_ of Arg ^14^; (iii) measured free energies of hydration of uncharged variants of charged residues; and (iv) a collection of 72 different computed and experimentally derived estimates of 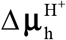, the proton free energy of hydration at 298 K.

**Figure 1:**
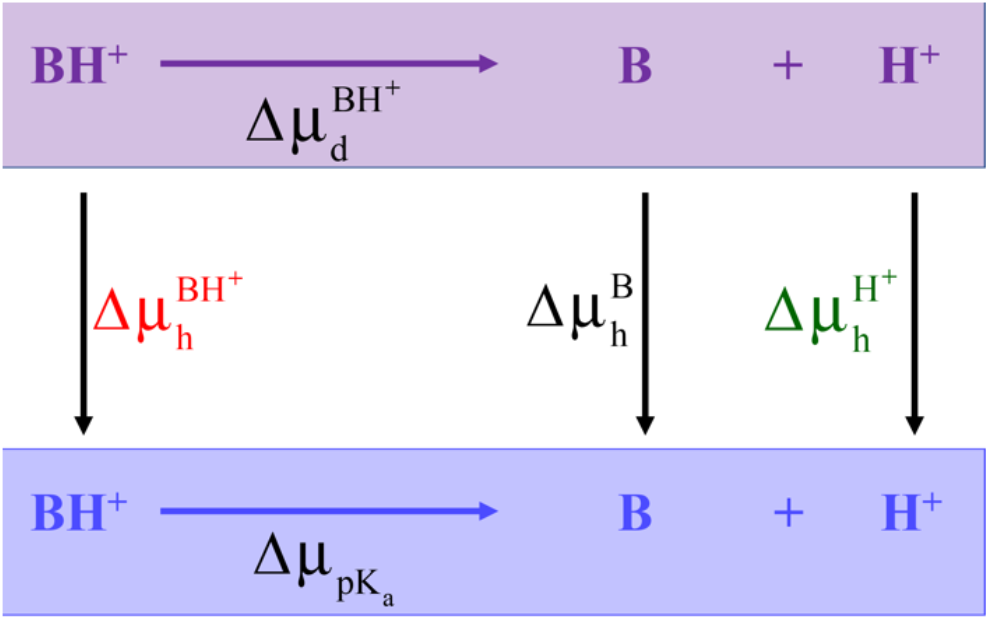
Illustration of the TCPD approach. The schematic shows the transfer of a strong base and the proton dissociation reaction from the gas phase (purple) into the aqueous phase (blue).

The TCPD approach is motivated by the separate albeit complementary efforts of Sitkoff et al.,^15^ Pliego and Riveros^16^ and Zhang et al.^17^. We follow closely the approach of Pliego and Riveros who estimated absolute values for Δµ_h_ for 30 different univalent ions, many of which are organic ions. They relied on the estimates for the free energy of hydration of the proton provided by Tissandier et al., ^18^. There were challenges with obtaining Δµ_h_ for the model compound that mimics the Arg^+^ sidechain because of persistent uncertainties regarding its pK_a_^14^ in the aqueous phase and the absence of data for proton dissociation in the gas phase. Improved estimates are now available for all the relevant quantities. We adapted these for obtaining TCPD derived values for the free energies of hydration of charged amino acids as illustrated in **Figure 1**. The TCPD derived estimates serve as updated reference values for the free energies of hydration of charged amino acids, which will depend on the choice one makes for 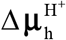. We show that free energies calculated using the polarizable AMOEBA (Atomic Multipole Optimized Energetics for Biomolecular Applications) forcefield for water and model compound mimics of the sidechains of Arg^+^, Lys^+^, Asp^-^, and Glu^-^ reproduce the trends obtained using the TCPD analysis. The simulations were then analyzed to obtain comparative assessments of hydration structures. This yields a physical picture for the trends we observe for free energies of hydration of model compounds that mimic sidechains of charged residues.

## 2. METHODS

### 2.1 Details of the TCPD approach

Concepts underlying the TCPD approach are summarized in **Figure 1**. The model compounds used as mimics for the ionized versions of the sidechains are 1-propylguanidinium (Arg^+^), 1-butylammonium (Lys^+^), acetate (Asp^-^), and propionate (Glu^-^). For bases, deprotonation reactions in the gas and aqueous phases are written as 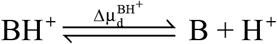 and 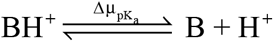, respectively. Here, 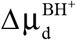 quantifies the change in free energy that accompanies the dissociation of a proton from the base in the gas phase ^19^ whereas 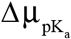 is the equivalent quantity in the aqueous phase ^20^. The free energy of proton dissociation in the aqueous phase can be obtained from knowledge of the pK_a_ for the model compound of interest whereby 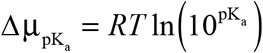. Here, *R =* 1.98717 × 10^−3^ kcal /mol-K and *T* is set to 298 K. This approach is the complement of the method used by Jorgensen and Briggs ^21^. Using measured values of 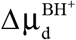 and of 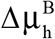 combined with estimation of 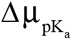 based on measured pK_a_ values, and knowledge of the free energy of hydration of the proton 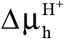 allows the usage of TCPD analysis to obtain the free energy of hydration 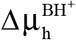 of the protonated base as shown in Equation (1):

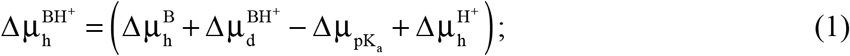

Likewise, for acids, the free energies of hydration (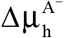) of the deprotonated forms are calculated using Equation (2):

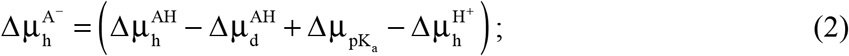

Here, 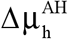 is the free energy of hydration of the protonated form of the acid, which is measured directly ^22^, 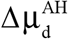 quantifies the change in free energy that accompanies the dissociation of a proton from the acid in the gas phase; the value to be used for 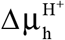 is identical to that of Equation (1). In Equations (1) and (2), 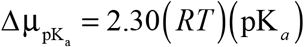 is estimated using measured pK_a_ values of the base ^23^ or acid, respectively.

The TCPD approach has been proposed ^24^ and used in the literature ^16^, and its usage requires accurate measurements of the relevant parameters ^17^. These are now available in the form of accurate proton dissociation / association energies in the gas phase, well-established and reliable values for 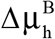 and 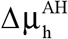, and improved estimates of the pK_a_ values, specifically for the Arg^+^ sidechain. As for 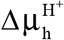, we found 72 distinct estimates (**Table 1**) in the literature. The mean value is -260.89 ± 5.82 kcal / mol. In this work, we obtain TCPD estimates for the free energies of hydration of Arg^+^, Lys^+^, Asp^-^, and Glu^-^ as a function of 72 distinct estimates for the free energy of hydration of the proton.

**Table 1.** 72 distinct values for the proton free energy of hydration collated from the literature. Rows that are bold-faced are estimates from analysis of experimental data without the use of any simulations. This is noteworthy because the mean value of -260.89 is closest to estimates derived from parsing of experimental data, sans any calculations.

| $\Delta\mu_h^{H^+}$ (kcal/mol) | Reference | Methods |
| --- | --- | --- |
| -247.00 | Lamoureux et al.,<sup>25</sup> | Molecular dynamics simulations with a polarizable forcefield based on the Drude model |
| -251.43 | Schmid et al.,<sup>26</sup> | Hydration entropy is obtained based on the thermodynamics of the dissociation of water. Hydration enthalpy is obtained based on the relation between hydration entropy and hydration enthalpy proposed by Krestov<sup>27</sup>. |
| -252.39, -253.08, -253.18, -253.27, -253.30, -253.49, -253.54, -253.66, -254.90, -255.23 | Marković et al.,<sup>28</sup> | Quantum mechanical (QM) calculations with the solvation model based on density (SMD) |
| -258.26 | Grossfield et al.,<sup>11</sup> and this work | The intrinsic free energy of hydration was estimated by Grossfield et al., using the known free energy difference between K<sup>+</sup> and H<sup>+</sup> and AMOEBA derived intrinsic free energies of hydration for K<sup>+</sup>. The value reported by Grossfield et al., was -252.5 kcal / mol for the intrinsic free energy of hydration. Beck has estimated the Galvani potential for the AMOEBA water model to be -0.25 V, which corresponds to a correction of -5.76 kcal / mol, leading to the final estimate of -258.26 kcal/mol for the corrected free energy of hydration of the proton for the AMOEBA model. |
| -254.60 | Asthagiri et al.,<sup>29</sup> | Quasi-chemical theory |
| -253.40 | Latimer et al.,<sup>30</sup> | The Born equation with additional assumptions is used to calculate the free energy of cations and ions. |
| -253.40 | Carvalho and Pliego<sup>31</sup> | Cluster continuum quasi-chemical theory |
| -254.28 | Marcus<sup>32</sup> | A correction term for the compression of the space available to the ion on its transfer from its gaseous to its aqueous standard states is made to the value -252.39 kcal/mol (-1056 kJ/mol) |
| -256.93 | Duignan et al.,<sup>33</sup> | Estimates are made using the established difference in the free energy of hydration between Li<sup>+</sup> and H<sup>+</sup>. The free energy of hydration of Li<sup>+</sup> is calculated using DFT interaction potentials with molecular dynamics simulations (DFT-MD) combined with a modified version of the quasi-chemical theory. |
| -259.50 | Pearson<sup>13</sup> | **Based on the absolute potential of hydrogen electrode** |
| -260.28 | Vlcek et al.,<sup>34</sup> | A correction for the surface potential is made to the value from cluster pair approximation. |
| -260.50 | Friedman and Krishnan<sup>35</sup> | **Parsing of data using a reference salt tetraphenyl arsonium tetraphenyl borate (TATB) method** |
| -260.76 | Fawcett<sup>36</sup> | **Fit to data from measurements of the ionic work function** |
| -258.80 | Yu et al.,<sup>37</sup> | Simulations based on a forcefield that uses the Drude model for atomic polarizabilities |
| -262.40 | Zhan and Dixon<sup>38</sup> | QM calculations for the ion-water cluster and a self-consistent reaction field model for the interaction between the cluster and solvent |
| -262.38, -261.86 | Hofer and Hünenberger<sup>39</sup> | QM/MM simulations and thermodynamic integration |
| -261.73, -262.23, -262.27, -262.67, | Tawa et al.,<sup>40</sup> | QM calculations for the ion-water cluster and a self-consistent reaction field model for the interaction between the cluster and solvent |
| -261.86, -262.38, -262.89 | Prasetyo et al.,<sup>41</sup> | QM/MM simulations and thermodynamics integration |
| -262.91 | Reif and Hünenberger<sup>42</sup> | Inferred from hydration structures obtained using classical molecular dynamics simulations |
| -263.79 | Tuttle et al.,<sup>43</sup> | Cluster pair approximation method |
| -263.98 | Tissandier et al.,<sup>18</sup> | Cluster pair approximation method |
| -264.20 | Pollard and Beck<sup>44</sup> | Quasi-chemical theory analysis of cluster pair approximation |
| -256.75, -254.26, -259.75, -262.46, -254.75, -261.50, -252.05, -268.35, -265.22, -265.15, -267.54, -267.30, -266.67, -265.17, -266.04, -265.43, -265.46 | Matsui et al.,<sup>45</sup> | Based on the relationship between pKa value, free energy of solvation of the neutral and charged versions of small molecules and free energy of solvation of proton. Experimental pKa values are used. Free energy of solvation of the neutral and charged versions of small molecules are calculated from QM with continuum solvent model. |
| -266.10, -268.40, -266.60, -265.10, -267.80, -265.80, -264.40, -271.30, -267.70, -263.70, -269.30, -266.80, -266.10, -268.00 | Kelly et al.,<sup>46</sup> | Cluster pair approximation method |
| -266.40 | Rossini and Knapp <sup>47</sup> | Uses quantum chemical density functional theory calculations of proton affinity in the gas phase and use of the Poisson equation to compute solvation contributions. |
| -252.50, -266.70 | Bryantsev et al., <sup>48</sup> | Cluster Continuum Model |
| -267.88 | Ishikawa and Nakai <sup>49</sup> | Cluster Continuum Model |

### 2.2 Set up of simulations using the AMOEBA forcefield

The free energy calculations we report here are an extension of the recent simulations performed by Zeng et al., ^4^. The authors developed parameters of the requisite model compounds for the AMOEBA forcefield. The details of the forcefield parameterization, set up of the simulations, and free energy calculations may be found in the work of Zeng et al., ^4^. In the interest of completeness, we include a summary of the overall simulation setup and analysis of simulation results.

Simulations were performed using the TINKER-OpenMM package ^50^. For each model compound, the simulations were performed using a cubic water box with periodic boundary conditions. The initial dimensions of the central cell were set to be 30×30×30 Å^3^. All molecular dynamics simulations were performed using the RESPA integrator ^51^ with an inner time step of 0.25 ps and an outer time step of 2.0 fs in isothermal-isobaric ensemble (NPT) ensemble. The target temperature was set to be one of 275 K, 298 K, 323 K, 348 K, or 373 K and the target pressure was set to be 1 bar. The temperature and pressure were controlled using a stochastic velocity rescaling thermostat ^52^ and a Monte Carlo constant pressure algorithm ^53^, respectively. The particle mesh Ewald (PME) method ^54^ with PME-GRID being 36×36×36, and *B*-spline interpolation ^55^, with a real space cutoff of 7 Å was used to compute long-range corrections to electrostatic interactions. The cutoff for van der Waals interactions was set to be 12 Å. This combination has been verified ^56^ in previous work for AMOEBA-based free energy simulations ^57^.

### 2.3 Free energy calculations

We used the Bennett Acceptance Ratio (BAR)^58^ and Multistate Bennett Acceptance Ratio (MBAR)^59^ methods to estimate the intrinsic free energies of hydration (Δµ_h,intrinsic_) for the model compounds of interest. Details of the simulation setup are identical to those of Zeng et al., ^4^.The solute is grown in using two different Kirkwood coupling parameters λ_vdW_ and λ_el_ that scale the strengths of solute-solute and solute-solvent van der Waals and electrostatic interactions. A series of independent molecular dynamics simulations were performed in the NPT ensemble for different combinations of λ_vdW_ and λ_el_. A soft-core modification of the Buffered-14-7 function was used to scale the van der Waals interactions as implemented in Tinker-OpenMM ^50^. For each pair of λ values, we performed simulations, each of length 6 ns, at the desired temperature and a pressure of 1 bar. We then used the TINKER bar program and the pymbar package https://github.com/choderalab/pymbar to calculate the free energy difference between neighboring windows defined in terms of the scaling coefficients. For every combination of λ_vdW_ and λ_el_, we set aside the first 1 ns simulation as part of the equilibration process. In Appendix A, we show results from our analysis of the BAR-derived free energy estimates for different λ schedules.

### 2.4 From intrinsic free energies of hydration to corrected values

Following the rigorous definitions of free energies of hydration ^7^, it follows that the transfer of an ionic solute from a fixed position in the gas phase (vacuum) to a fixed position in the water sets up a contribution from crossing of the interface between the gas and aqueous phases ^60^. This interface cannot be captured in simulations that use periodic boundary conditions ^61^. Accordingly, the free energies of hydration that we obtain using protocols described in sections 2.2 and 2.3 are intrinsic free energies. These have to be corrected by the contributions of the surface potential, known as the Galvani potential and denoted as Φ_G_. The corrected free energy of hydration is calculated using the relation: Δµ_h,corrected_ = Δµ_h,intrinsic_ + *q*Φ_G_^62^.

Beck has estimated the Galvani potential for the AMOEBA water model to be -0.25 V / *e* ^63^. This translates to -5.76 kcal / mol / *e*. Accordingly, the corrected free energies of hydration, for the AMOEBA forcefield, are estimated using the intrinsic free energies of hydration calculated as described in section 2.3, Beck’s estimate for the Galvani potential, and setting *q* to +1 for Arg^+^ and Lys^+^ and *q* to -1 for Asp^-^ and Glu^-^.

## 3. RESULTS

### 3.1 Free energies of hydration calculated using the TCPD approach

Values of free energies of hydration for model compound mimics of Arg^+^, Lys^+^, Asp^-^, and Glu^-^ sidechains were calculated at 298 K using the TCPD approach – see Equations (1) and (2) and **Figure 1**. The difference between the gas phase basicity ^64^ and 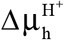 quantifies the relative importance of bond energy and the free energy of solvation. Positive values for this difference imply that the favorable free energy of hydration of the proton cannot compensate for the loss of bond energy in the gas phase. In order to achieve the target pK_a_ value for the ionizable moiety, suitably large magnitudes for 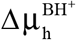 and 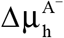 help offset the loss of bond energy in the gas phase. Values of 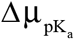 are derived from measurements of pK_a_ values for the relevant model compound mimics of Arg^+^, Lys^+^, Asp^-^, and Glu^-^. The pK_a_ values of all four model compounds were taken from the Physical/Chemical Property Database (PHYSPROP)^3^ database and they reflect updates from the measurements of Fitch et al., ^14^ and Xu et al., ^65^. These measurements move the consensus estimate for the pK_a_ of Arg up from 12.6 and 13.2^15^ to 13.6. Values for free energies of hydration for uncharged constructs, i.e., 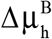 and 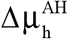 were obtained from the Hydration Free Energy Database curated by Mobley and Guthrie ^7^.

The gas phase dissociation energies 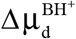 and 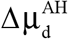, estimated from gas-phase basicity measurements ^64^, are available from the literature for three of the four model compounds. Gas phase basicity measurements of sidechain mimicking model compounds were taken from the National Institute of Standards and Technology (NIST) for acetic acid, propanoic acid, and 1-butylammonium. Experimental data for 1-propylguanidine are unavailable. Instead, we used results from gas phase quantum-mechanical calculations for 1-methylguanidine ^64^. These calculations yield excellent agreement for gas phase basicities as compared to experimental values obtained for a range of model compounds. The differences in electronic structure between 1-propylguanidine and 1-methylguanidine are considerably smaller than the differences in electronic structures of 1-methylguanidine and guanidine. Accordingly, we use the calculated gas phase basicity value for 1-methylguanidine as a more suitable proxy for the basicity of 1-propylguanidine. This is relevant because that in their deployment of the TCPD approach, Zhang et al.,^17^ used guanidine as a model compound to mimic the Arg sidechain. The difference in gas phase basicities of 1-methylguanidine and guanidine is greater than 7 kcal / mol. Accordingly, the use of basicity values for guanidine results in a significant overestimation of the magnitude of the of Δµ_h_ for the Arg^+^ sidechain.

The free energy of hydration of the proton 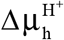 is a crucial parameter that determines the outputs we obtain from the TCPD approach. We combed the literature and found at least 72 distinct estimates for 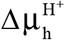 – see **Table 1** and **Figure 2**. As summarized in **Table 1**, the approaches used to obtain estimates of 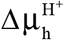, combine quantum-mechanical calculations, empirical considerations / prescriptions, and bespoke interpretations of experimental data for whole salts or pK_a_ values. The distribution of tabulated values yields a mean of -260.89 ± 5.82 kcal / mol for 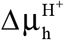 at 298 K. Instead of choosing a specific value for 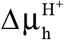, we compute the values of the free energies of hydration (Δµ_h_) for Arg^+^, Lys^+^, Asp^-^, and Glu^-^ for each of the 72 tabulated values of 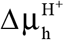. The values we obtain for Δµ_h_ are plotted as a function of values used for 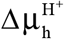 (**Figure 3**).

**Figure 2:**
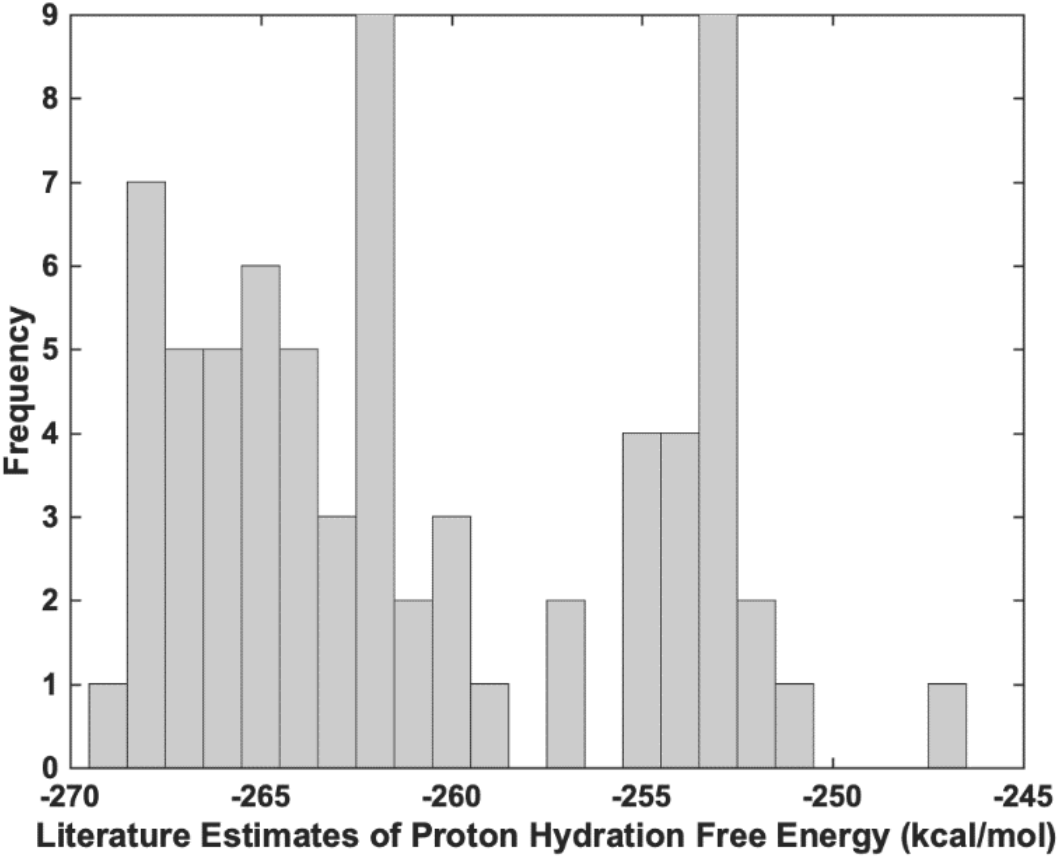
Distribution of tabulated values for the proton hydration free energy at 298 K. These values are listed in **Table 1** and were collated from the literature.

**Figure 3:**
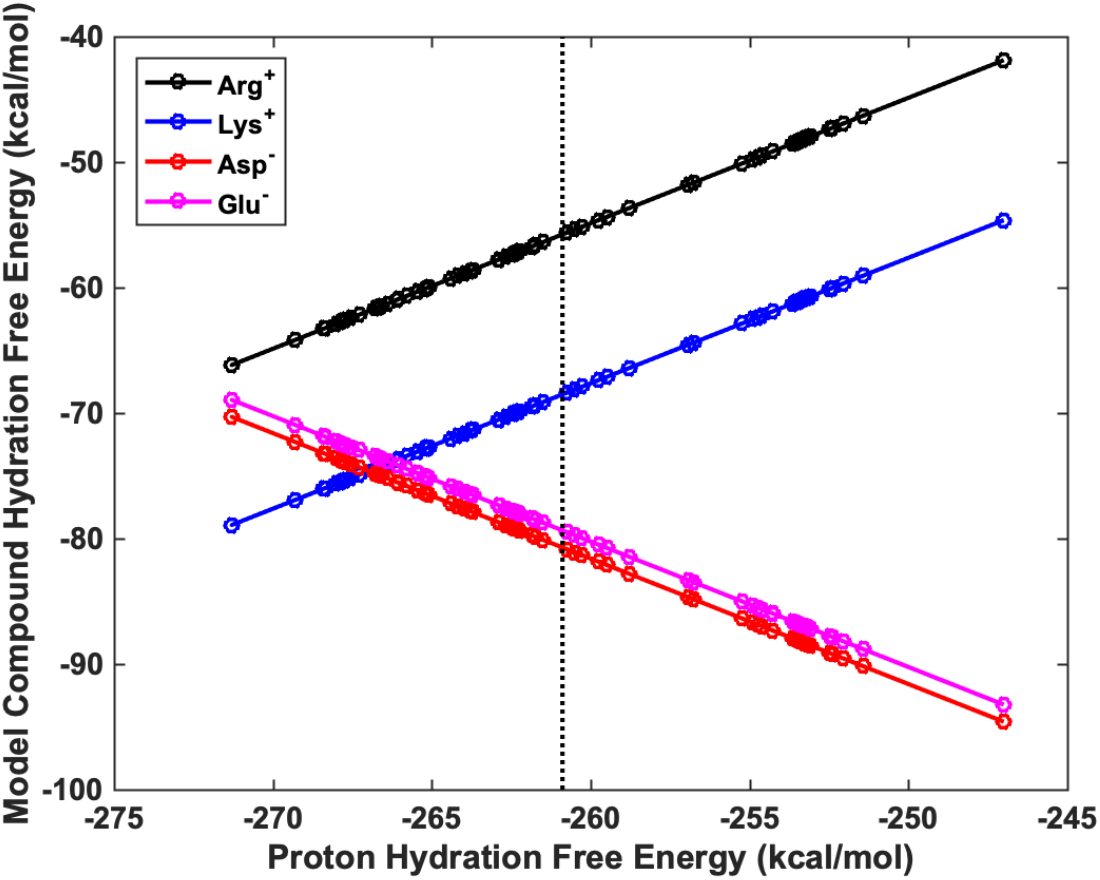
TCPD derived free energies of hydration at 298 K. The data are plotted against the literature derived proton hydration free energies (circles). The solid lines join the circles and are included as guides to the eye. The vertical dotted line intersects the abscissa at the mean value of -260.89 kcal / mol for the proton hydration free energy.

Results from application of the TCPD approach are summarized in **Table 2**. In addition to the values of 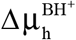 and 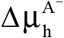 obtained using the mean value of 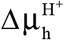 from **Table 1**, we also tabulate the values used for measured and / or calculated gas phase basicity values, measured pK_a_ values, and free energies of hydration for the uncharged forms of the model compounds. These estimates are for a temperature of 298 K. As expected, the estimated free energies of hydration are large and negative. However, the TCPD based estimates revealed unexpected trends. Despite being a strong base, the estimated value Δµ_h_ is ∼12 kcal / mol less favorable for Arg^+^ when compared to Lys^+^. Further, the estimates for Δµ_h_ of Arg^+^ and Lys^+^ are smaller in magnitude than those for Asp^-^ and Glu^-^. The acids are more favorably hydrated than bases – an observation that is concordant with results for free energies of hydration of small anions versus cations ^66^.

**Table 2:**
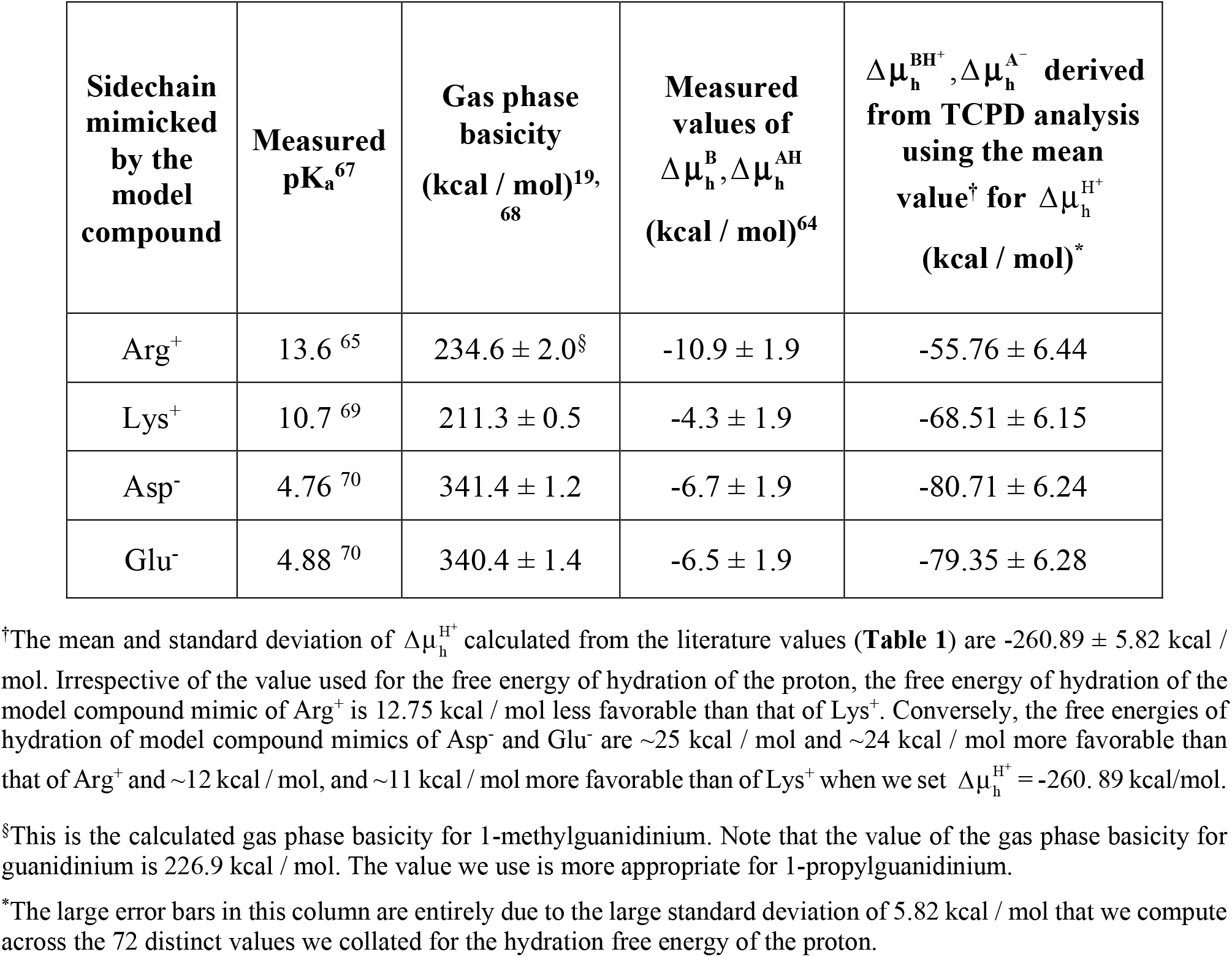
**Summary of inputs to and outputs from the TCPD approach used for estimating values free energies of hydration for Arg**^**+**^, **Lys**^**+**^, **Asp**^**-**^, **and Glu**^**-**^.

### 3.2. Prescription for comparing computed free energies of hydration to TSPD estimates

The large and persistent uncertainties in estimates of the free energy of hydration of the proton make it impossible to obtain precise, experimentally derived values of Δµ_h_ for Arg^+^, Lys^+^, Asp^-^ and Glu^-^. However, one can prescribe a measure of consistency that can be used to judge the accuracy of a forcefield calculation. If we denote the forcefield and water model specific proton free energy as 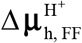, then the forcefield derived estimates of Δµ_h_ for Arg^+^, Lys^+^, Asp^-^ and Glu^-^ would have to be similar to the TCPD derived estimate obtained by setting 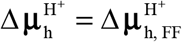. This level of consistency is the best one can hope for pending the availability of a data-driven consensus regarding the precise value for 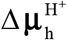. The approach we propose for assessing consistency with the TCPD approach guards against imposing false standards based on definitive assertions that in reality will always depend on the choice one makes for 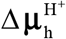.

### 3.3 Free energy calculations based on the AMOEBA forcefield yield values that are consistent with a proton free energy of hydration of -258.26 kcal / mol

For each of the model compounds, we used the AMOEBA forcefield and water model ^71^ used to calculate intrinsic free energies of hydration at four different temperatures *viz*., 275 K, 298 K, 323 K, and 348 K. The results are summarized in **Table 3**.

**Table 3:** **Summary of results obtained from calculations of intrinsic free energies of hydration (Δµ**_**h**,**intrinsic**_**) derived from free energy calculations using the AMOEBA forcefield**.

| Sidechain mimicked by the model compound | $\Delta\mu_{h,intrinsic}$ (kcal / mol) 275 K | $\Delta\mu_{h,intrinsic}$ (kcal / mol) 298 K | $\Delta\mu_{h,intrinsic}$ (kcal / mol) 323 K | $\Delta\mu_{h,intrinsic}$ (kcal / mol) 348 K |
| --- | --- | --- | --- | --- |
| Arg<sup>+</sup> | -47.63 ± 0.11 | -46.72 ± 0.11 | -45.67 ± 0.11 | -45.45 ± 0.09 |
| Lys<sup>+</sup> | -61.22 ± 0.10 | -60.49 ± 0.09 | -59.53 ± 0.09 | -58.99 ± 0.08 |
| Asp<sup>-</sup> | -90.39 ± 0.08 | -89.91 ± 0.08 | -89.02 ± 0.07 | -88.15 ± 0.07 |
| Glu<sup>-</sup> | -86.84 ± 0.12 | -86.16 ± 0.12 | -85.24 ± 0.12 | -84.32 ± 0.11 |

Combining estimates for the Galvani potential in the AMOEBA water model^63^ and the intrinsic proton free energy of hydration reported by Grossfield et al.,^11^ the corrected value for the free energy of hydration of the proton at 298 K is -258.26 kcal / mol for the AMOEBA forcefield. We calculate the root mean squared deviation (RMSD) to be 0.97 kcal / mol between the corrected free energies of hydration calculated using the AMOEBA forcefield and those estimated using the TCPD approach by setting 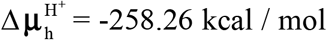 (**Table 4**). The RMSD being within a kcal / mol suggests that the free energies of hydration we obtain for Arg^+^, Lys^+^, Asp^-^ and Glu^-^ are consistent with experimental data for the experimentally derived TCPD based estimates if we use the proton free energy of hydration that is consistent with that of the AMOEBA forcefield.

**Table 4:**
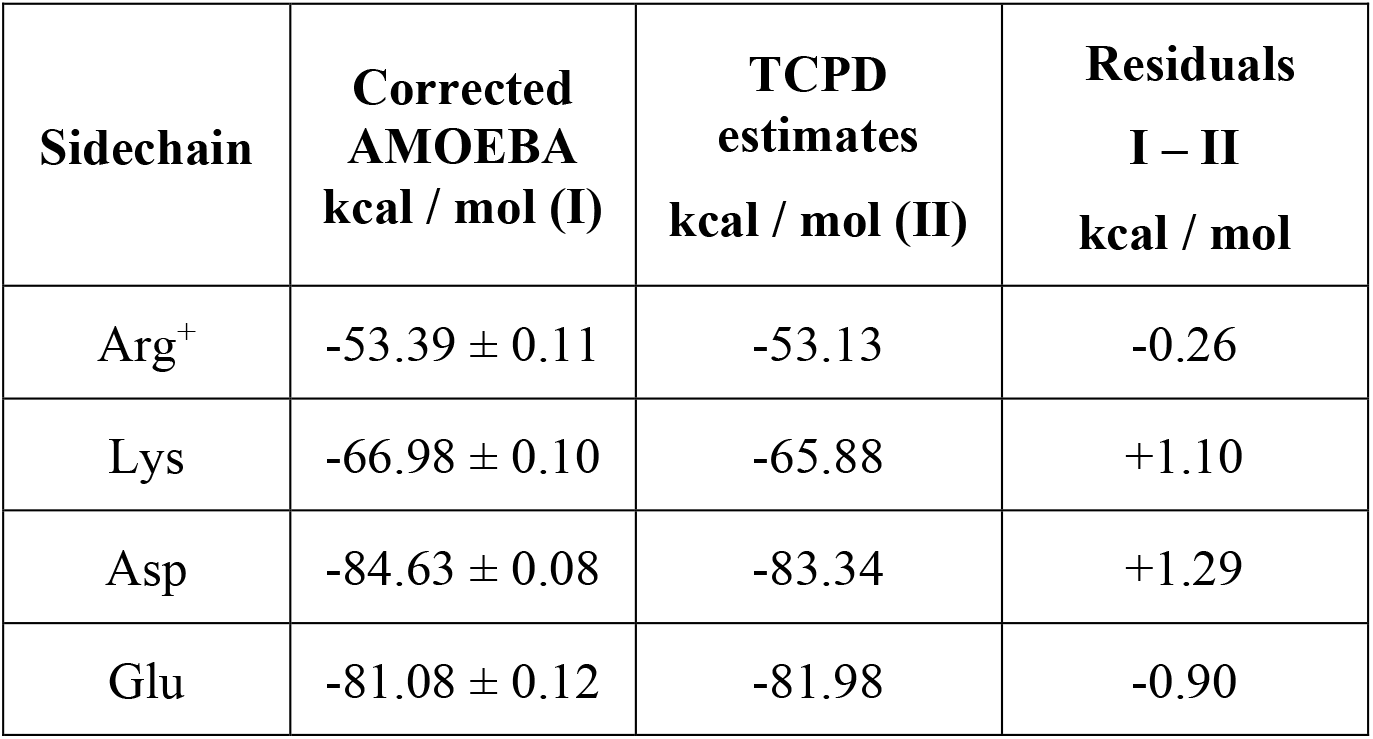
**Comparison of corrected free energies of hydration from AMOEBA at 298 K to TCPD estimates obtained using a proton free energy of hydration of -258**.**26 kcal / mol**.

| Sidechain | Corrected<br>AMOEBA<br>kcal / mol (I) | TCPD<br>estimates<br>kcal / mol (II) | Residuals<br>I – II<br>kcal / mol |
| --- | --- | --- | --- |
| Arg<sup>+</sup> | -53.39 ± 0.11 | -53.13 | -0.26 |
| Lys | -66.98 ± 0.10 | -65.88 | +1.10 |
| Asp | -84.63 ± 0.08 | -83.34 | +1.29 |
| Glu | -81.08 ± 0.12 | -81.98 | -0.90 |

*How do the corrected estimates for* Δ*µ*_*h*_ *obtained using simulations based on the AMOEBA forcefield compare to estimates obtained using other instantiations of polarizable forcefields*? To answer this question, we compared our results to those reported by Lin et al., using the classical Drude oscillator model ^72^. The molecular ions studied by Lin et al., include 1-methylguanidinium and acetate. Lin et al., report a value of -84.7 ± 0.1 kcal / mol for acetate. In comparison, the corrected value we obtain for acetate using the AMOEBA forcefield is -84.63 ± 0.08 kcal / mol. Further, as noted in Table 4, the value we obtain using the AMOEBA forcefield is with 1.29 kcal / mol of the value we derive from the TCPD approach, providing we set the hydration free energy of the proton to be -258.26 kcal / mol – the value for the AMOEBA forcefield. Lin et al., reported a value of -59.3 kcal / mol for 1-methylguanidinium. The value we obtain from AMOEBA simulations for 1-propylguanidinium is -53.39 kcal / mol. Although, direct comparisons between the free energies of hydration for the model compounds are confounded by differences in sidechain structure, Table 4 clearly shows that the TCPD derived estimate, which uses the gas phase basicity for 1-methylguanidinium, is closer to the AMOEBA derived value for 1-propylguanidinium. If we use the mean value of -260.89 for the free energy of hydration of the proton, we estimate the free energy of hydration for 1-propylguanidinium to be -55.76 kcal / mol as shown in Table 2. This is closer to the value we derive using simulations based on the AMOEBA forcefield when compared to the value reported by Lin et al., for 1-methylguanidinium.

### 3.4 Insights from analysis of the temperature dependence of calculated free energies of hydration

**Table 3** shows how the intrinsic free energies of hydration vary with temperature for each of the model compounds. The consistent trend is of the intrinsic free energies of hydration becoming less favorable as temperature increases. We fit the temperature dependent data for the intrinsic free energies of hydration to the integral of the Gibbs-Helmholtz equation in order to estimate the intrinsic enthalpy of hydration (Δ*h*) and intrinsic heat capacity of hydration (Δ*c*_P_) at a reference temperature of 298 K. In doing so, we assume that values of Δ*h* and Δ*c*_P_ are independent of temperature, a conjecture that is supported by the linear increase in the magnitudes of 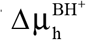 and 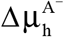 with increasing temperature. The integral of the Gibbs-Helmholtz equation is written as:

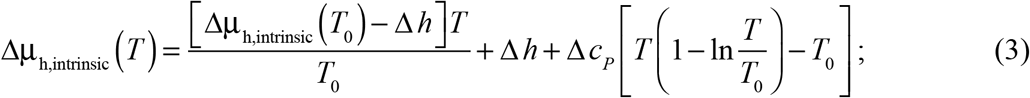

To use Equation (3), we set *T*_0_ = 298 K, substitute the calculated value of Δµ_h,intrinsic_(*T*_0_), and estimate Δ*h* and Δ*c*_P_ using a Levenberg-Marquardt nonlinear least squares algorithm. The values we obtain for Δ*h* and Δ*c*_P_ are shown in **Table 5**. As a test of the quality of the fit, we compare the values of Δµ_h,intrinsic_(*T*) from free energy calculations to those obtained using Equation (3). For the latter, we use the parameters listed in **Table 5**. The comparisons are shown in **Figure 4**.

**Table 5:** **Parameters for Δ*h* and Δ*c***_**P**_ **extracted from non-linear least squares analysis of computed temperature dependent free energies and fits to Equation 3**.

| Sidechain mimicked by the model compound | Estimated $\Delta h$ at 298 K (kcal / mol) | Estimated $\Delta c_p$ (cal / mol-K) |
| --- | --- | --- |
| Arg<sup>+</sup> | -57.24 | 69.39 |
| Lys<sup>+</sup> | -70.37 | 29.98 |
| Asp<sup>-</sup> | -98.65 | -44.97 |
| Glu<sup>-</sup> | -96.62 | -8.75 |

**Figure 4:**
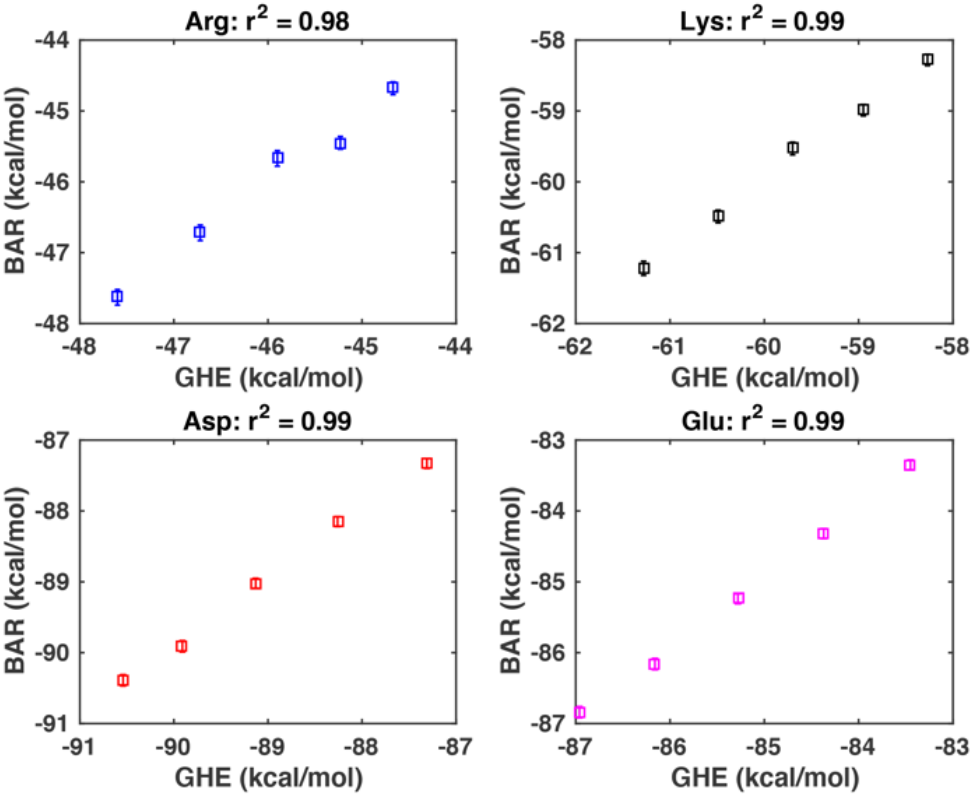
**Assessment of the correlation between temperature-dependent intrinsic free energies of hydration calculated using the integral of the Gibbs-Helmholtz equation (GHE) and direct calculations from AMOEBA-based simulations.**

### 3.5 Analysis of temperature dependent intrinsic values of enthalpy and entropy of hydration

Using the Gibbs-Helmholtz equation and parameters shown in **Table 5**, we estimated the temperature dependence of the intrinsic enthalpy and entropy of hydration using Equation (4) below:

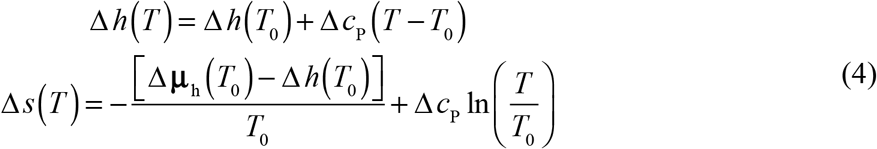

The results are shown in **Figure 5**. There is a clear difference in the temperature dependencies for basic versus acidic molecules. With increasing temperature, the enthalpy of hydration becomes less favorable for Arg^+^ and Lys^+^ while it becomes more favorable for Asp^-^ and Glu^-^. The unfavorable entropic contribution to the free energy of hydration stays roughly constant for Lys^+^ and decreases with increasing temperature for Arg^+^. In contrast, the unfavorable entropic contribution to the free energy of hydration increases with increasing temperature for Asp^-^ and Glu^-^. Each of the results shown in **Figure 5** is a direct consequence of the negative heat capacity of hydration for Asp^-^ and Glu^-^, which contrasts with the positive heat capacity of hydration for Arg^+^ and Lys^+^ and all other model compounds that mimic backbone and sidechain moieties.

**Figure 5:**
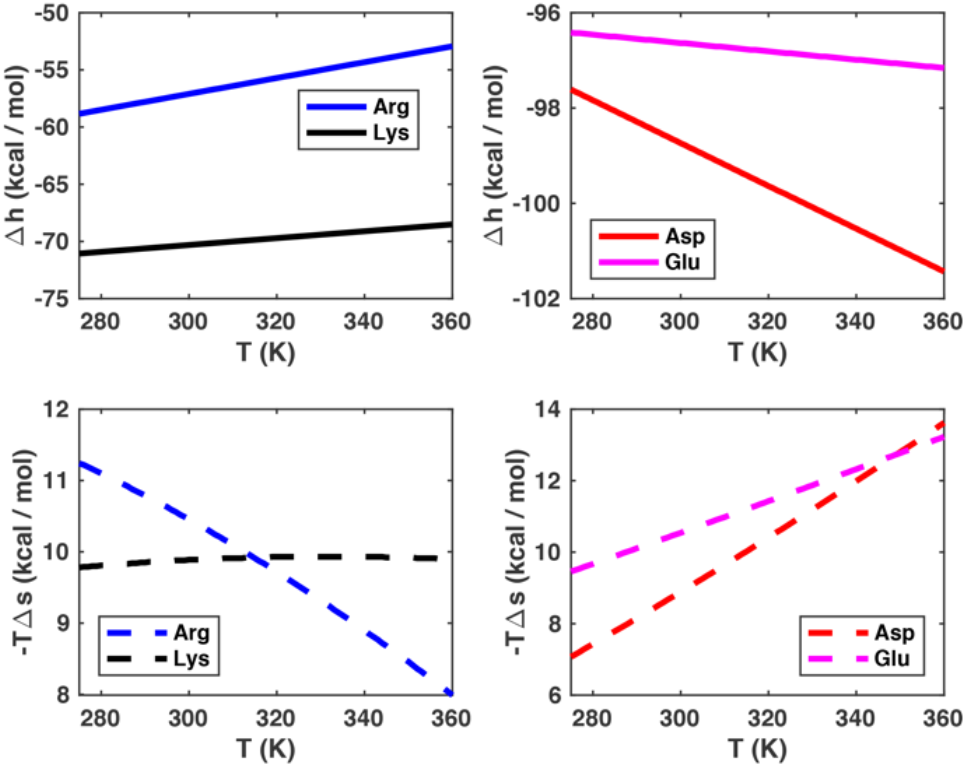
Temperature dependent enthalpies and entropies of hydration for the four model compound mimics of Arg^+^, Lys^+^, Asp^-^, and Glu^-^. These decompositions were calculated using the integral of the Gibbs-Helmholtz equation and parameters from **Table 5**.

To understand the origins of the observations summarized in **Figure 5**, we performed three sets of reference simulations, one for the Cl^−^ ion and two for alchemic variants of the Cl^−^ ion. The anionic Cl^−^ ion is weakly polarizable in the AMOEBA model and carries a net charge of -*e*. We computed free energies of hydration for the following temperatures: 275 K, 298 K, 323 K, 348 K, and 373 K. The results for intrinsic free energies of hydration as a function of temperature are shown in **Table 6**. Here, we also show results for two alchemic versions of the Cl^−^ ion namely, an uncharged version Cl^0^ and a cationic version Cl^+^, where we flip the sign of the charge. From the temperature dependent values of the intrinsic free energies of hydration for Cl^−^, Cl^0^, and Cl^+^ we extract estimates for the intrinsic enthalpy of hydration Δ*h* at 298 K and the heat capacity of hydration Δ*c*_P_. These values are also tabulated in **Table 6**. Comparison of the parameters in **Tables 5** and **6** reveal the following: Anions that mimic Asp^-^ and Glu^-^ and the Cl^−^ ion have negative Δ*c*_P_ values. The magnitude of Δ*c*_P_ decreases and approaches zero as the length of the alkyl chain increases – see comparisons of Δ*c*_P_ values for mimics of Asp^-^ versus Glu^-^. The Δ*c*_P_ values are positive for model compound mimics of Arg^+^, Lys^+^, and the alchemic Cl^+^ ion. The magnitude of the positive Δ*c*_P_ increases with hydrophobicity and surprisingly, 1-propylguanidinium has a higher Δ*c*_P_ when compared to the neutral, alchemic Cl^0^ solute. These numerical findings prompted detailed comparisons of hydration structures, and these are presented next.

**Table 6:** **Parameters for Δ*h* and Δ*c***_**P**_ **extracted from non-linear least squares analysis of computed temperature dependent free energies and fits to Equation 3**.

| Solute | $\Delta\mu_h$ at 298 K | Estimated $\Delta h$<br>at 298 K<br>(kcal / mol) | Estimated $\Delta c_P$<br>(cal / mol-K) |
| --- | --- | --- | --- |
| $\text{Cl}^-$ | $-86.17 \pm 0.06$ | -87.72 | -46.55 |
| $\text{Cl}^0$ | $2.50 \pm 0.04$ | -2.70 | 63.31 |
| $\text{Cl}^+$ | $-65.30 \pm 0.05$ | -68.65 | 11.35 |

### 3.6 Comparative analysis of hydration structures around the different solutes

Molecular theories for hydrophobic and hydrophilic hydration rest on comparative analyses of hydration structures around solutes and the effects of solutes on density inhomogeneities within water. The chemical structures of the mimics of Arg^+^, Lys^+^, Asp^-^, and Glu^-^ are subtly or significantly different from another. We computed the spatial density profiles of water molecules around each of the solutes. The results from this analysis are summarized pictorially in **Figure 6**. Here, each panel shows regions around each solute where the density of oxygen and hydrogen atoms from water molecules rises above a prescribed cutoff value – see caption. These calculations emphasize the accumulation of water molecules around the functional groups within each solute.

**Figure 6:**
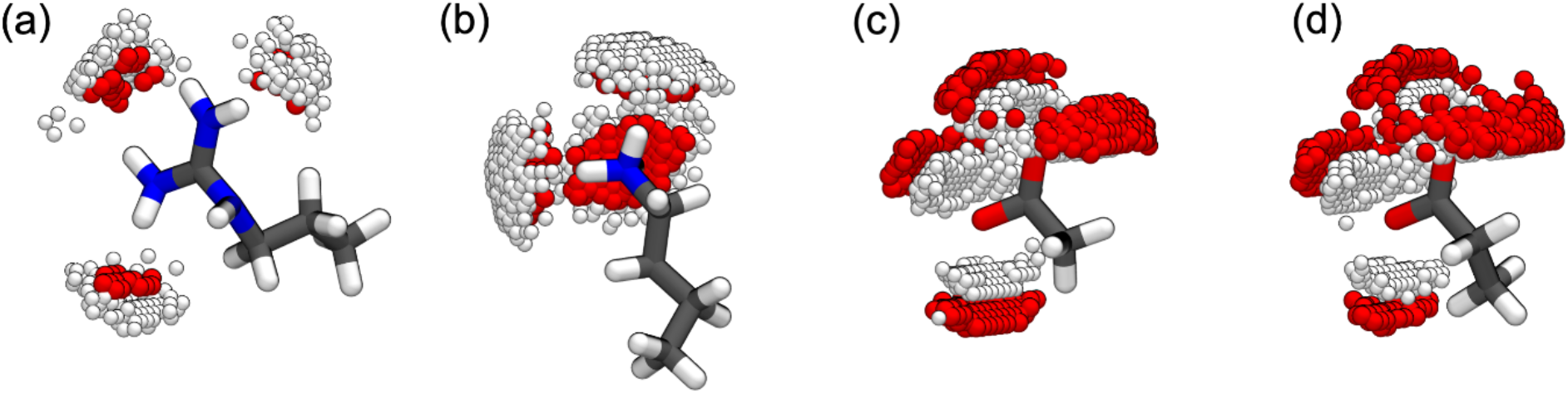
Hydration structures around the model compounds mimicking the sidechains of (a) Arg^+^, (b) Lys^+^,(c) Asp^-^ and (d) Glu^-^. The red and white spheres around the model compounds denote areas with a time-averaged density of water oxygen and hydrogen atoms being larger than 0.2 Å^-3^. Positions further than 2Å away from the model compound are not shown. To calculate the density, we define two vectors 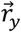 and 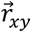 for the model compounds to align all the frames in the trajectory. All the coordinates for atoms in the frame are translated and rotated so that the central atom in the model compound is in the origin of the simulation box and 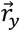 points to the *y* direction and 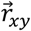 is in the *x-y* plane. For (a), the central atom is the carbon atom in the guanidine group, 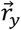 is the vector pointing from the central atom to the nitrogen atom bonded with two carbon atoms, 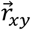 is the vector pointing from the central atom to one of the nitrogen atoms bonded with two hydrogens. For (b), the central atom is the nitrogen atom, 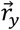 is the vector pointing from the central atom to the carbon atom bonded with nitrogen, 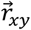 is the vector pointing from the central atom to one of the three hydrogens bonded with it. For (c) and (d), the central atom is the carbon atom bonded with oxygen atoms, 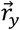 is the vector pointing from the central atom to the carbon atoms bonded with the central atom, 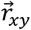 is the vector pointing from the central atom to one of the oxygen atoms. Each panel was made using VMD ^73^.

The distinction between hydrophobic and hydrophilic hydration is typically attributed to differences in hydration structure within the first hydration shell ^74^, to the spatial organization of the first shell with respect to the bulk ^74-75^, to density inhomogeneities in the vicinity of the solute ^76^, and their long-range effects ^77^. We quantify hydration structures in terms of two-parameter probability distribution functions. Here, we follow the approach of Gallagher and Sharp ^75^ and compute the joint radial and angular distribution functions ρ(*r*,θ) where the definition of *r* and θ are as shown in **Figure 5**. For neat water, the distribution functions are computed using all pairs of water molecules that are within 8 Å of one another. For solute-solvent systems, water molecules within the first hydration shell around each solute were used to compute the ρ(*r*,θ) distributions. For each system, ρ(*r*,θ)δ*r*δθ quantifies the probability that a pair of water molecules will be in a distance interval *r* and *r*+δ*r and* have relative orientations that are between θ and θ+δθ.

Optimal hydrogen bonding is realized for short distances and values of θ that are close to zero. These results are shown in **Figure 8**. The basin corresponding to *r* ≈ 2.8 Å and values of θ < 10° are evident in each of the four panels of **Figure 8** and these peaks represent the optimal hydrogen bonded geometries for water molecules. This peak becomes pronounced for Cl^0^ and Cl^+^, which are the alchemic neutral and cationic solutes, respectively. The density in the interval 3 Å < *r* < 7 Å *and* 0° < θ < 60°is significantly higher for water in the presence of the anion when compared to neat water or in the presence of the uncharged, non-polar Cl^0^ solute. In this region, there are two distinct peaks, and this increased density is consistent with the calculated negative Δ*c*_P_ values in that it points to distinct structural preferences of water molecules in the presence of anionic solutes. In contrast, for water around the cationic Cl^+^ solute, the density is dispersed across the interval 3 Å < *r* < 7 Å and 40° < θ < 120°i.e., reflected about an axis that intersects the ordinate at ≈ 40°. As with the uncharged Cl^0^ solute, there is a sharp density for water in the regime corresponding to optimal hydrogen bonding and this is weakened as *r* increases for the Cl^+^ solute.

**Figure 7:**
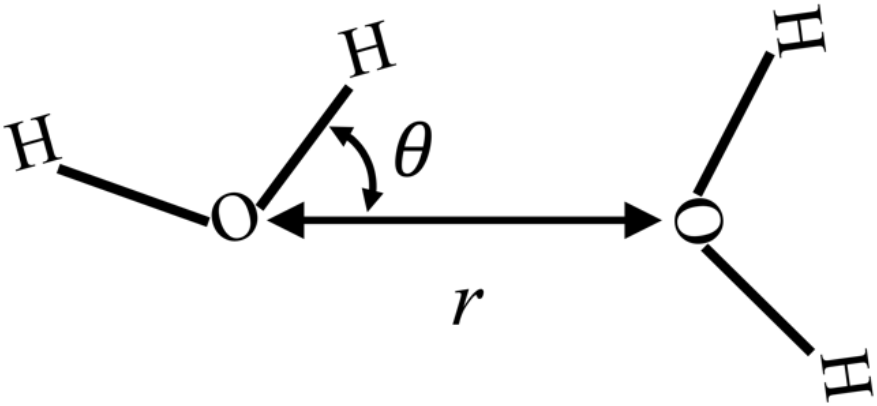
Definition of *r* and θ for characterizing water structures as defined by Gallagher and Sharp ^75^. Here, *r* is the distance between two water oxygen atoms, θ is the smallest angle in the four O-O_x_-H_x_ angles, where H_x_ is bonded with O_x_.

**Figure 8:**
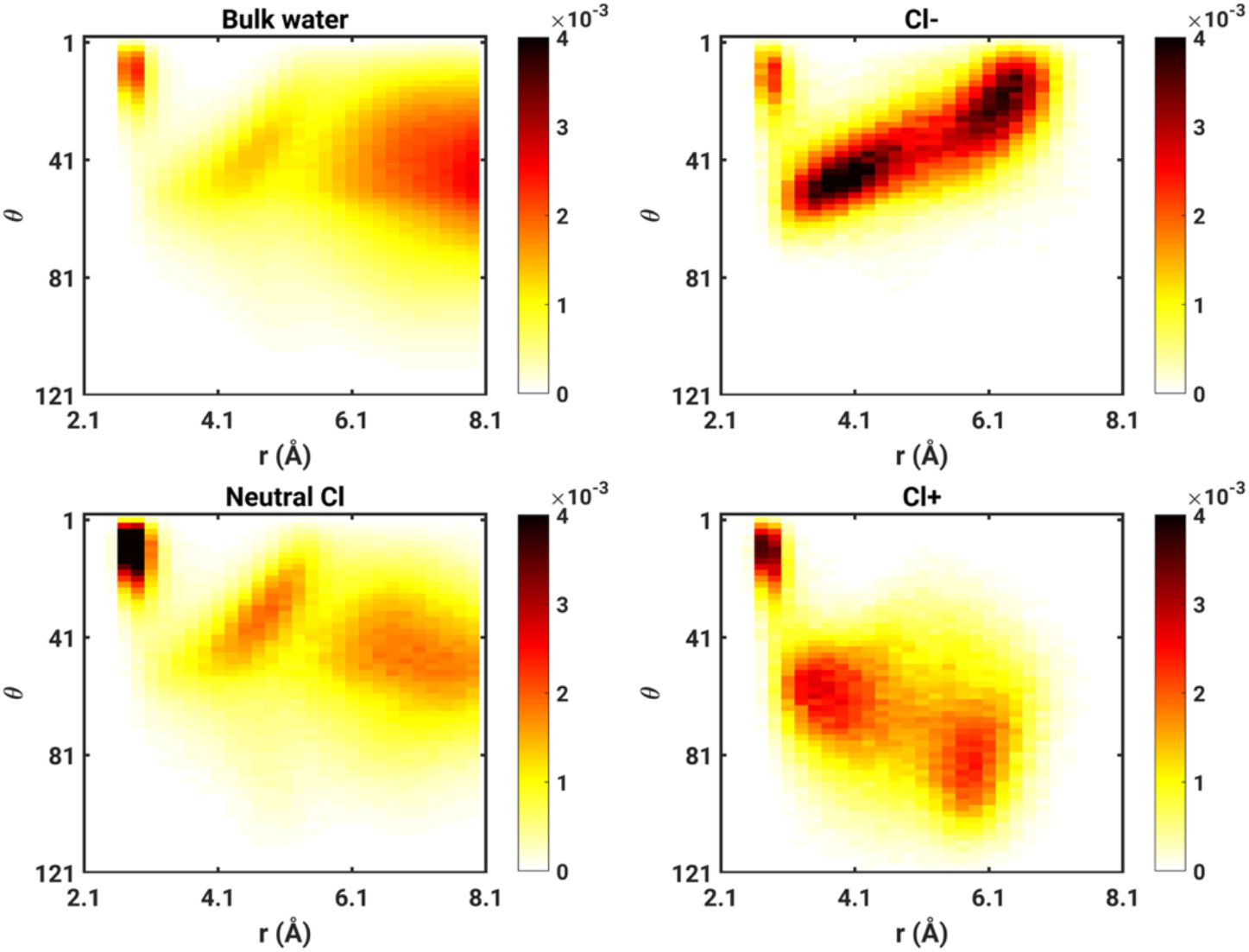
Joint distributions *r* and θ (see Figure 6) for bulk water and the waters in the first solvation shell of Cl^-^, Cl^0^ (neutral Cl), and Cl^+^ at 298K. For Cl-, Cl+ and the neutral Cl, the water is considered in the first solvation shell if the distance from the solute to the water oxygen atom is smaller than the radius of the 1^st^ solvation shell, which are 4.0 Å, 3.9 Å and 5.4 Å for Cl-, Cl+ and neutral Cl, respectively. For each system, the histogram is calculated from a 6 ns long trajectory with a saving interval of 1 ps. The bin size is 0.2 Å for r and 2° for θ.

**Figure 9** shows difference density distributions Δρ(*r*,θ), where for each solute X, the difference distribution is calculated as ρ_X_(*r*,θ) – ρ_w_(*r*,θ), where ρ_X_(*r*,θ) and ρ_w_(*r*,θ) are the joint distributions for the solute-solvent system with solute X and neat water, respectively. In these difference distribution plots, regions where there is an enhancement of density vis-à-vis neat water are in hot colors, whereas the regions where there is a depletion of density compared to neat water are shown in cool colors. The difference distributions highlight the fundamental differences between hydrophobic hydration seen for Cl^0^ and the anion versus cation systems.

**Figure 9:**
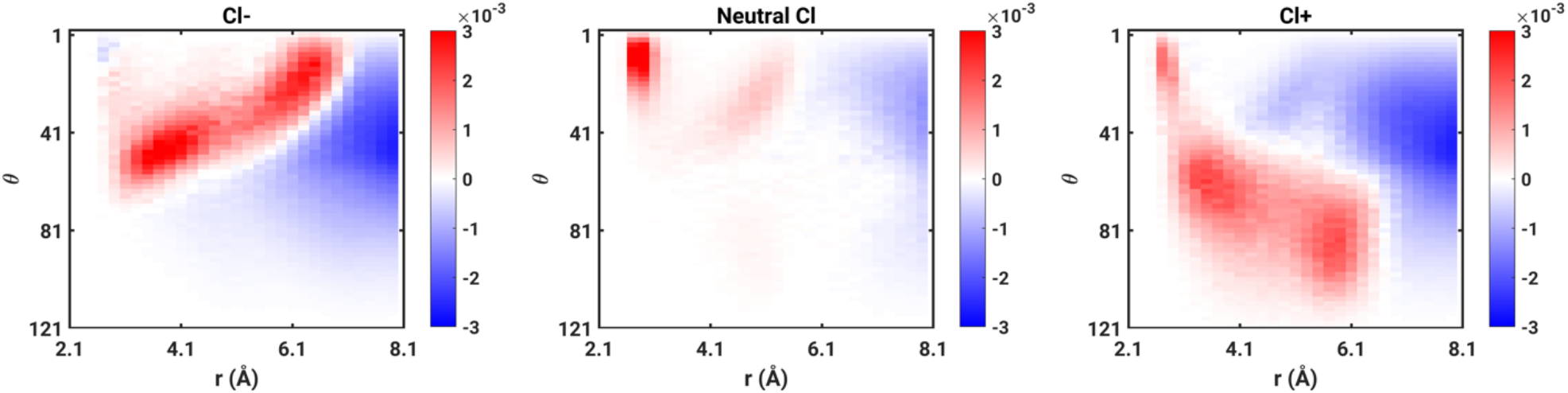
The difference density distributions for the three reference systems.

Next, we analyzed the difference density distributions for model compounds that mimic Arg^+^, Lys^+^, Asp^-^, and Glu^-^, respectively. The results, shown in **Figure 10**, show that the difference density distributions for Arg^+^ and Lys^+^ are clearly very different from one another. When compared to neat water, there is a significant increase of density in the basin corresponding to *r* ≈ 2.8 Å and θ < 10° for Arg^+^. This increase is similar to that of the model hydrophobic solute Cl^0^. For Lys^+^, the density in the basin corresponding to *r* ≈ 2.8 Å and values of θ < 10°is considerably lower than that of neat water or the model hydrophobic solute Cl^0^. Instead, there is a pronounced increase in density in the region corresponding to 3 Å < *r* < 7 Å and 40° < θ < 120°, which is concordant with the observations for Cl^+^, although the distribution is considerably more uniform for Lys^+^. The difference density distributions for Asp^-^ and Glu^-^ are qualitatively similar to that observed for Cl^−^, showing increased preference for the interval 3 Å < *r* < 7 Å *and* 0° < θ < 60°and a clear weakening, vis-à-vis neat water, for the basin corresponding to *r* ≈ 2.8 Å and values of θ < 10°. Taken together, these features indicate that Arg^+^ behaves more like a hydrophobic solute when compared and Lys^+^, and the impact of the anionic moieties on water structure is mutually consistent, being qualitatively similar to that of Cl^−^ while also providing a rationalization for the negative heat capacities reported for these solutes.

**Figure 10:**
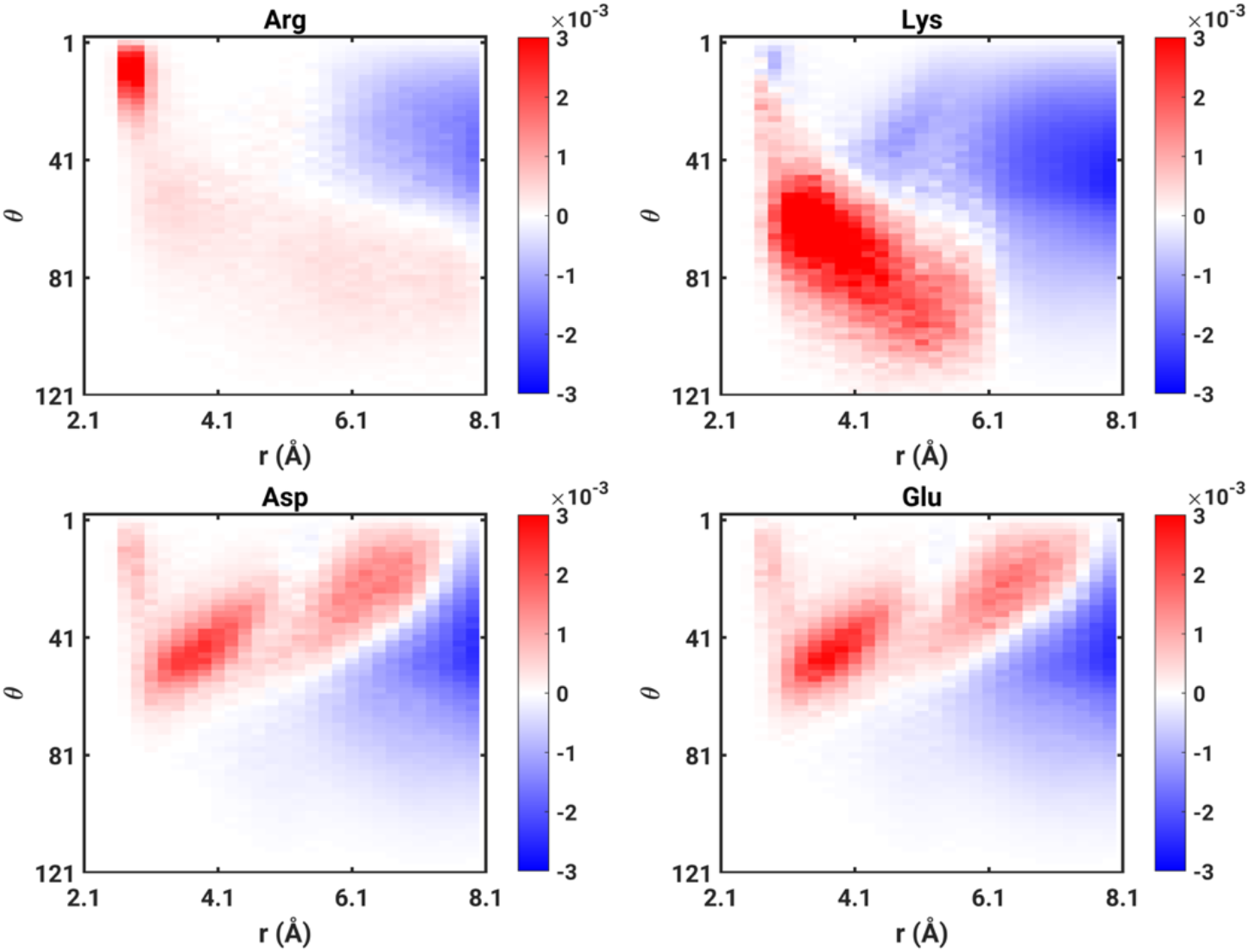
Difference density distributions for the model compounds mimicking the sidechains of the charged amino acids referenced to that neat water. The comparisons are shown for 298K.

Finally, we computed the numbers of water molecules that make up the first hydration shells around each of the four solutes. On average, there are 15 water molecules in the first shell around Arg^+^, ≈ 9 water molecules in the first shell around Asp^-^ and Glu^-^, and ≈ 5 water molecules in the first shell around Lys^+^. The larger numbers of water molecules around Arg are concordant with signatures of hydrophobic hydration when compared to Lys^+^, Asp^-^, or Glu^-^. Based on the estimates for the average numbers of water molecules in the first hydration shells, it follows that the free energy of hydration per water molecule is ≈ -3 kcal / mol for Arg^+^, ≈ -12 kcal / mol for Lys^+^, ≈ -10 kcal / mol for Asp^+^, and ≈ -9.6 kcal / mol for Glu^-^. These estimates suggest that the free energy cost for displacing individual water molecules from the first hydration shell will be smallest for Arg^+^, largest for Lys^+^, and ≈ 10 kcal / mol per molecule for Asp^-^ / Glu^-^.

## 4. DISCUSSION

### 4.1 Summary of main findings

We introduced our adaptation of the TCPD approach to estimate free energies of hydration from direct measurements of accessible quantities. The measured values are taken from the literature. Unfortunately, the persistent and large uncertainties associated with the free energy of hydration of the proton prevent us from obtaining precise values for Δµ_h_ of Arg^+^, Lys^+^, Asp^-^, and Glu^-^. However, the TCPD formalism allows one to estimate the relevant free energies of hydration that would be consistent with forcefield specific values for 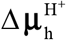. By collating 72 distinct values for 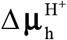 from the literature (**Table 1**), we were able to estimate mean values of the free energies of hydration for Arg^+^, Lys^+^, Asp^-^, and Glu^-^. (**Table 2**). Overall, the TCPD-based estimations point to clear trends regarding the free energies of hydration, and these are corroborated by direct calculations of intrinsic and corrected values obtained using the AMOEBA forcefield.

Using intrinsic free energies of hydration that were calculated at different temperatures, we obtained estimates for the enthalpy of hydration at a reference temperature of 298 K and the heat capacity of hydration. The latter was estimated by assuming that the heat capacity of hydration is independent of temperature. This is a reasonable assumption, and its validity is assessed by the quality of the agreement between the direct calculations of Δµ_h_ as a function of temperature and the values we obtain using the integral of the Gibbs-Helmholtz equation.

Overall, we report three main results: (1) Arg^+^ and Lys^+^ are non-equivalent in terms of their hydration preferences. Contrary to expectations based on partition coefficients between water and octanol ^78^, the hydration free energy we obtain for Arg^+^ is consistently less favorable than that of Lys^-^. Additionally, the heat capacity of hydration, which is positive for both species, is approximately 2.3 times larger for Arg^+^ when compared to Lys^+^. This large heat capacity of hydration is indicative of a more hydrophobic character for Arg. (2) The free energies of hydration for acidic residues are considerably more favorable than for basic residues. (3) The heat capacity of hydration is negative for Asp^-^ and Glu^-^. Negative heat capacities have been attributed to differences in hydration structure and the propagation of these effects beyond the first hydration shell. This is evident in the difference density distributions that we compute for the negatively charged solutes.

Measurements have reported negative heat capacities of hydration for whole salts, and this has been taken to imply that hydrophilic hydration, i.e., the hydration of anions as well as cations is associated with negative values for Δ*c*_P_^79^. This inference has been questioned by Sedlmeier and Netz ^77^. They showed that while the sum of heat capacities for anions and cations in whole salts can be negative, the negative Δ*c*_P_ values arise strictly from the anions. Their work uses the SPC/E water model and non-polarizable forcefields. Accordingly, it appears that the negative heat capacity of hydration for anions is a generic and robust attribute that does not depend on polarizability. Indeed, we find that the negative heat capacity is evident for Cl^−^ ions whereas the heat capacity is positive for alchemically transformed Cl^+^ and Cl^0^ solutes.

### 4.2 Relevance of our findings to recent studies on IDPs

Ongoing investigations of the determinants of driving forces of phase separation in IDPs and in RNA binding domains have revealed striking differences in the contributions of charged sidechains to the driving forces for phase separation ^80^. The most salient observation is that Arg and Lys are fundamentally different ^81^ as drivers of phase separation ^80, 82^. Replacing Arg residues with Lys significantly weakens the driving forces for phase separation in a variety of systems ^83^. In a similar vein, Sørensen and Kjaergaard recently showed that Arg-rich polyampholytic IDPs prefer considerably more compact conformations when compared to Lys-rich counterparts ^84^.

Various conjectures have been offered to explain the differences between Arg^+^ and Lys^+^ as drivers of phase separation and their differences on conformational equilibria of IDPs. The “Y-aromaticity” of Arg^+^, its high quadrupole moment, favorable interactions with pi systems, and the apparent ability to engage in water-mediated attractive interactions have been implicated as features that distinguish Arg^+^ from Lys^+^ residues as more potent drivers of phase separation ^2, 83^. However, a definitive rationale for explaining the differences between Arg and Lys and the manifest differences between the contributions of Asp / Glu versus Arg / Lys has been lacking. Our results suggest that differences in free energies of hydration and hydration structure are likely explanations for the intrinsic differences in charged sidechains as determinants of conformational and phase equilibria of IDPs – a hypothesis that is being tested in ongoing work.

## Appendix A

We assessed the statistical robustness of our of intrinsic free energies (Δµ_h,intrinsic_) by querying the sensitivity of our results obtained using BAR versus MBAR. We also assessed the impact of the lambda schedule on the estimates of estimates of Δµ_h,intrinsic_. **Table A1** shows comparative assessments of the temperature dependent values for Δµ_h,intrinsic_ obtained using BAR versus MBAR.

**Table A1.**
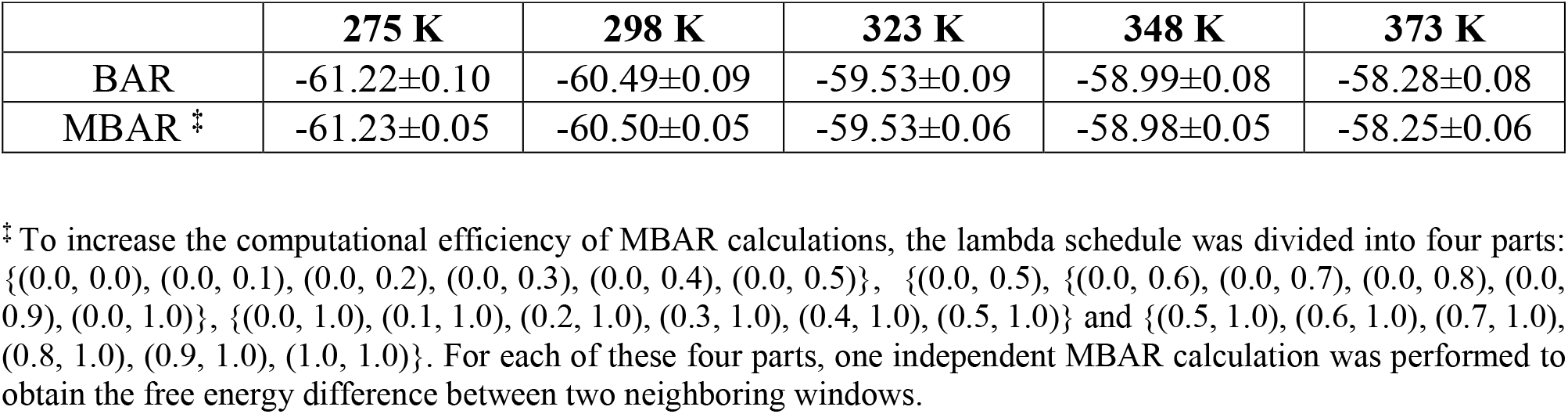
Intrinsic free energies of hydration of 1-butylammonium obtained at different temperatures calculated usiung BAR and MBAR methods. The lambda interval is 0.10 and the corresponding lambda schedule is listed in **Table A2**.

As summarized in **Table A1**, the values we obtain for the intrinsic free energies of 1-butylammonium are essentially identical obtained using BAR versus MBAR. This consistency prevails irrespective of the simulation temperature. Next, we assessed the impact of the lambda schedule on the free energy estimates obtained using BAR. The different lambda schedules used are tabulated in **Table A2**.

**Table A2.**
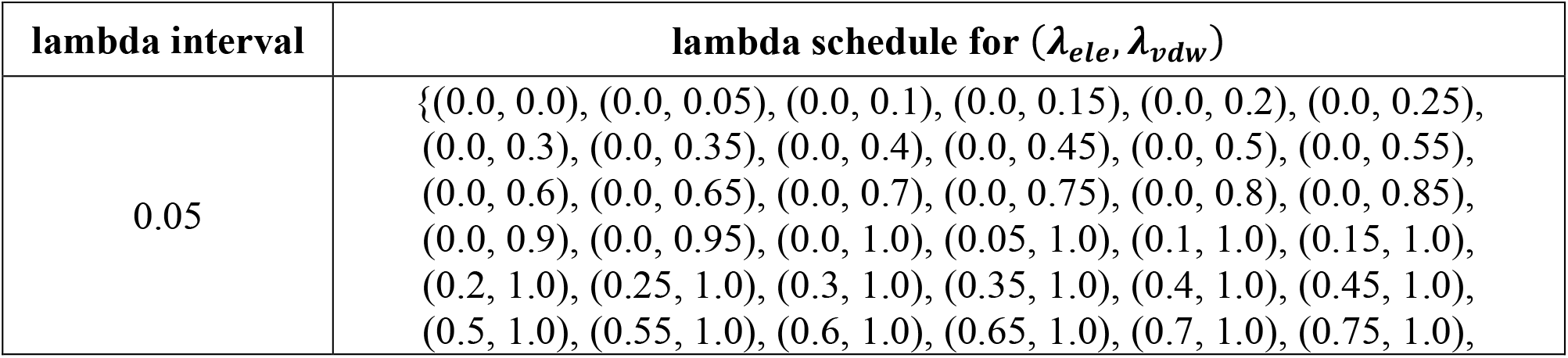
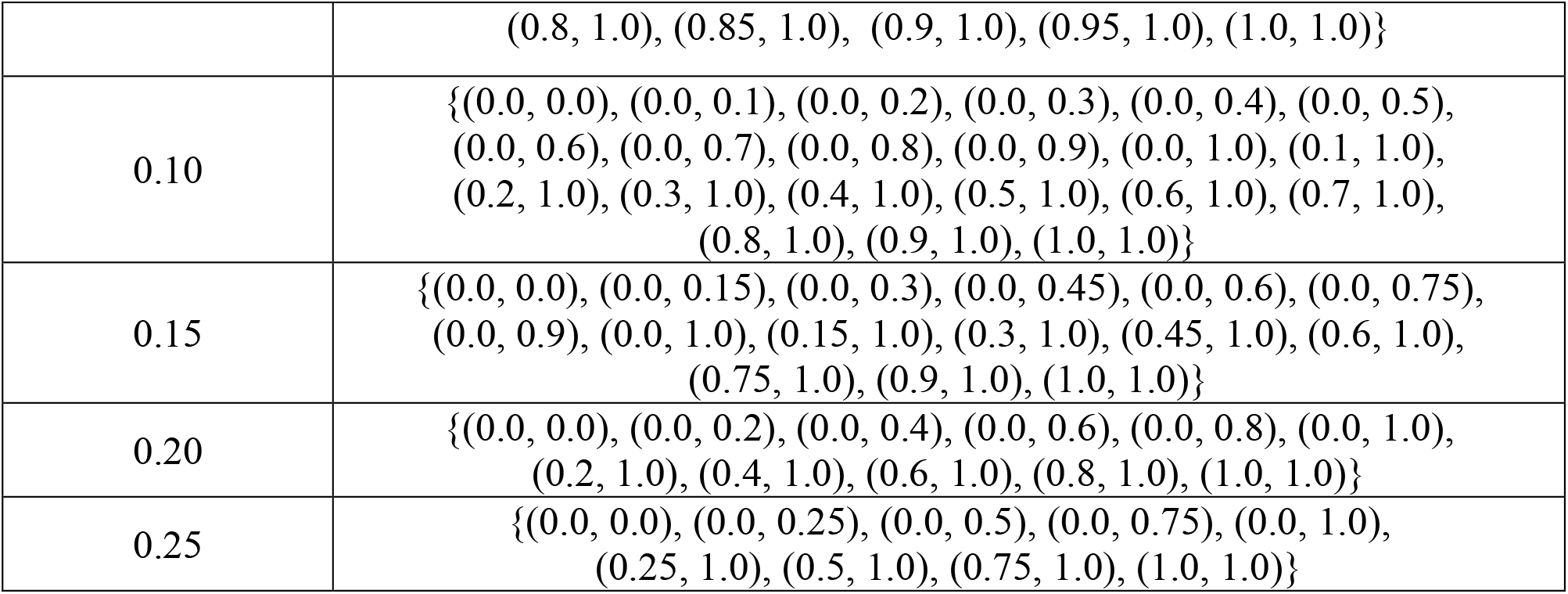
lambda schedules associated with distinct lambda intervals.

| lambda interval | lambda schedule for ($\lambda_{ele}, \lambda_{vdw}$) |
| --- | --- |
| 0.05 | {(0.0, 0.0), (0.0, 0.05), (0.0, 0.1), (0.0, 0.15), (0.0, 0.2), (0.0, 0.25), (0.0, 0.3), (0.0, 0.35), (0.0, 0.4), (0.0, 0.45), (0.0, 0.5), (0.0, 0.55), (0.0, 0.6), (0.0, 0.65), (0.0, 0.7), (0.0, 0.75), (0.0, 0.8), (0.0, 0.85), (0.0, 0.9), (0.0, 0.95), (0.0, 1.0), (0.05, 1.0), (0.1, 1.0), (0.15, 1.0), (0.2, 1.0), (0.25, 1.0), (0.3, 1.0), (0.35, 1.0), (0.4, 1.0), (0.45, 1.0), (0.5, 1.0), (0.55, 1.0), (0.6, 1.0), (0.65, 1.0), (0.7, 1.0), (0.75, 1.0), |
|  | (0.8, 1.0), (0.85, 1.0), (0.9, 1.0), (0.95, 1.0), (1.0, 1.0)} |
| 0.10 | {(0.0, 0.0), (0.0, 0.1), (0.0, 0.2), (0.0, 0.3), (0.0, 0.4), (0.0, 0.5), (0.0, 0.6), (0.0, 0.7), (0.0, 0.8), (0.0, 0.9), (0.0, 1.0), (0.1, 1.0), (0.2, 1.0), (0.3, 1.0), (0.4, 1.0), (0.5, 1.0), (0.6, 1.0), (0.7, 1.0), (0.8, 1.0), (0.9, 1.0), (1.0, 1.0)} |
| 0.15 | {(0.0, 0.0), (0.0, 0.15), (0.0, 0.3), (0.0, 0.45), (0.0, 0.6), (0.0, 0.75), (0.0, 0.9), (0.0, 1.0), (0.15, 1.0), (0.3, 1.0), (0.45, 1.0), (0.6, 1.0), (0.75, 1.0), (0.9, 1.0), (1.0, 1.0)} |
| 0.20 | {(0.0, 0.0), (0.0, 0.2), (0.0, 0.4), (0.0, 0.6), (0.0, 0.8), (0.0, 1.0), (0.2, 1.0), (0.4, 1.0), (0.6, 1.0), (0.8, 1.0), (1.0, 1.0)} |
| 0.25 | {(0.0, 0.0), (0.0, 0.25), (0.0, 0.5), (0.0, 0.75), (0.0, 1.0), (0.25, 1.0), (0.5, 1.0), (0.75, 1.0), (1.0, 1.0)} |

**Figure A1.**
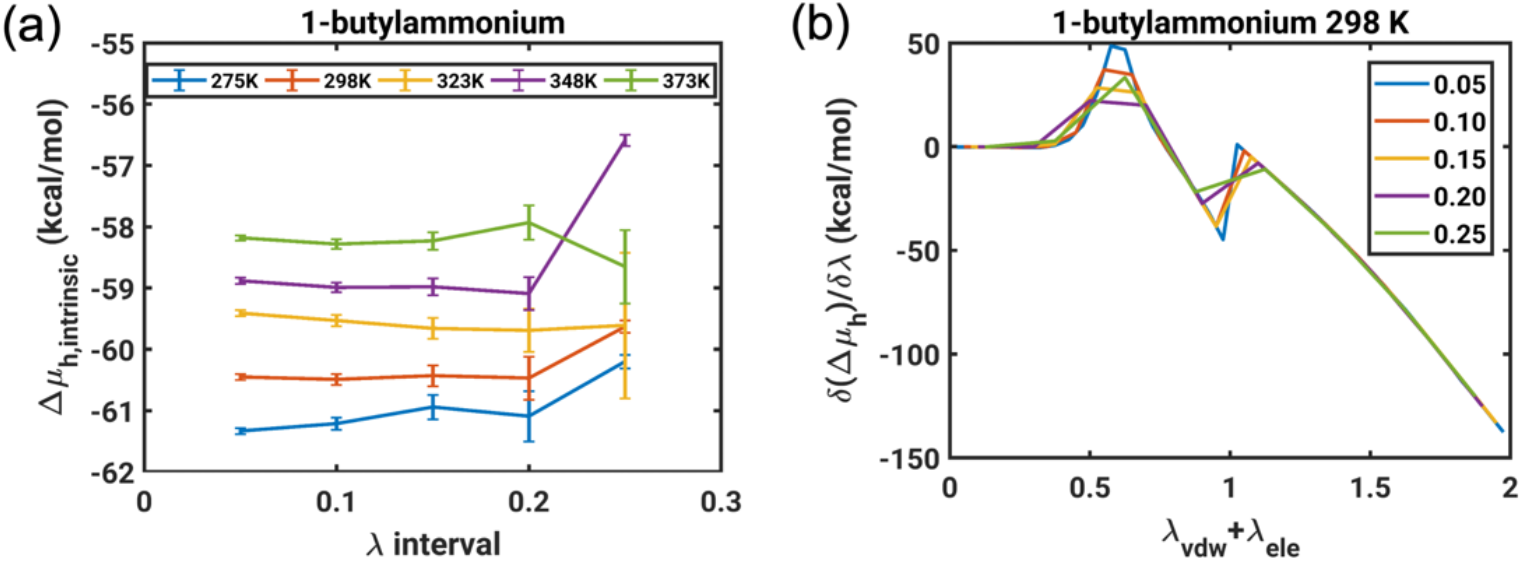
Assessment of the statistical robustness of free energy calculations based on the AMOEBA forcefield. **(a)** Intrinsic free energy of hydration (Δµ_h,intrinsic_) for 1-butylammonium at different temperatures calculated using the BAR method with different lambda intervals. The corresponding lambda schedules are listed in **Table A2. (b)** Plots of the derivative *δ*(Δ*μ*_*h*_)/ *δλ*for 1-butylammonium calculated using different lambda intervals using data obtained at 298 K.

The results we obtain for different lambda schedules are summarized in **Figure A1**. Notice that the estimates we obtain deviate significantly from one another only when the lambda schedule is too coarse i.e., for the lambda interval of 0.25. For more realistic and numerically conservative lambda schedules, the estimates we obtain are robust, and insensitive to the details of the lambda schedule.

## AUTHOR INFORMATION

The authors declare that there are no competing financial interests

## ACKNOWLEDGMENTS

We thank the anonymous reviewer for their constructive criticisms and helpful suggestions. We are grateful for the privilege of contributing this work as part of the festschrift celebrating Professor Dev Thirumalai’s numerous and influential contributions to physical chemistry, soft matter physics, and biophysics. Professor Thirumalai has been an inspiration and a generous mentor over the years. We are grateful to Chengwen Liu and Pengyu Ren for their collaboration on the original free energy calculations ^4^, developing parameters for the AMOEBA forcefield, and technical assistance. Our work is supported by grants from the US National Science Foundation (DMR1729783) and the US National Institutes of Health (5R01NS056114.

## Table of Content Graphic

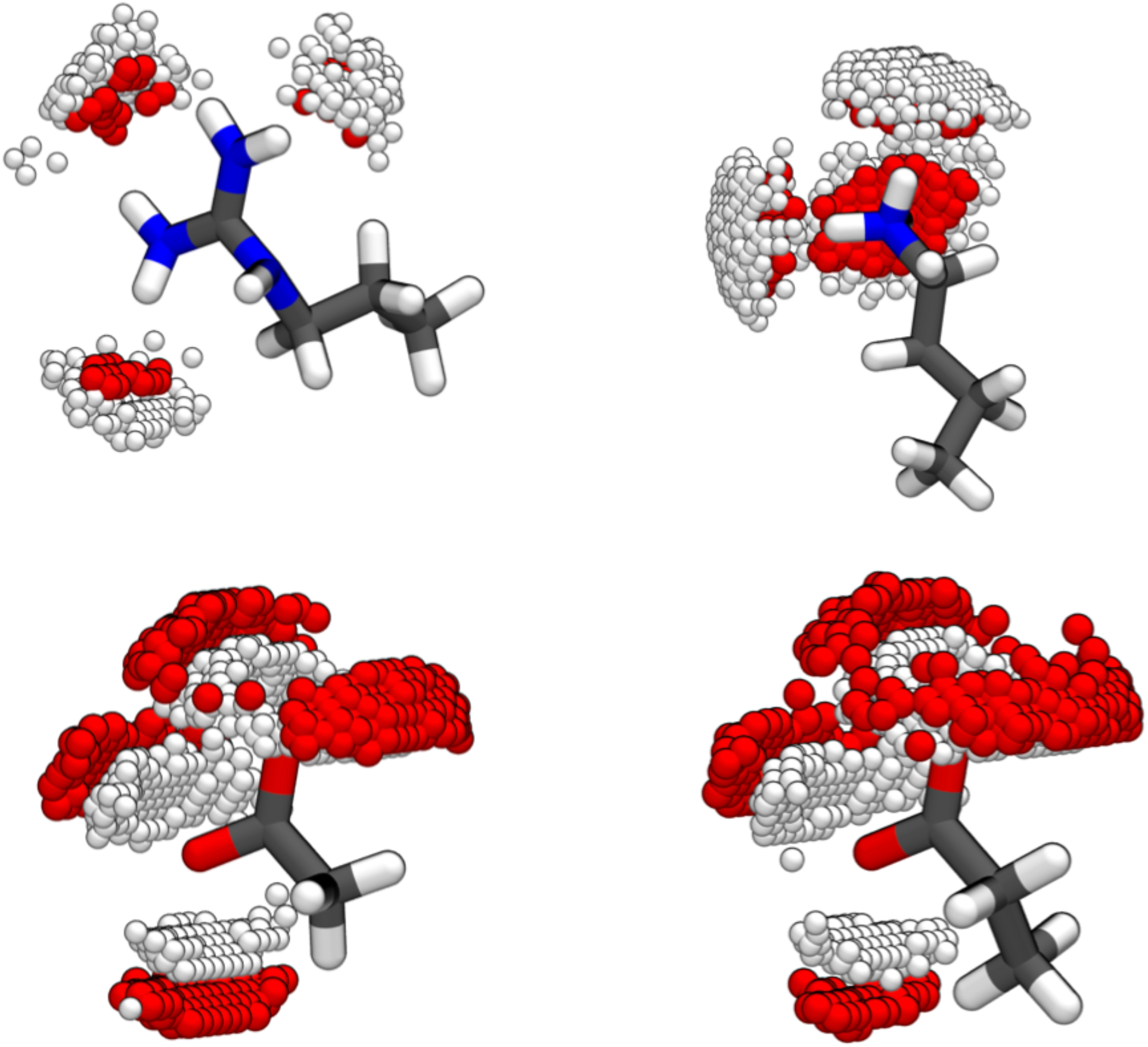

